# Impact of cell culture on the transcriptomic programs of primary and iPSC-derived human alveolar type 2 cells

**DOI:** 10.1101/2022.02.08.479591

**Authors:** Konstantinos-Dionysios Alysandratos, Carolina Garcia de Alba Rivas, Changfu Yao, Patrizia Pessina, Carlos Villacorta-Martin, Jessie Huang, Olivia T. Hix, Kasey Minakin, Bindu Konda, Barry R. Stripp, Carla F. Kim, Darrell N. Kotton

## Abstract

The alveolar epithelial type 2 cell (AEC2) is the facultative progenitor of lung alveoli tasked to maintain distal lung homeostasis. AEC2 dysfunction has been implicated in the pathogenesis of a number of pulmonary diseases, including idiopathic pulmonary fibrosis (IPF), highlighting the importance of human *in vitro* models of the alveolar epithelium. However, AEC2-like cells captured in cell culture have yet to be directly compared to their *in vivo* counterparts at single cell resolution. Here, we apply single cell RNA sequencing to perform head-to-head comparisons between the global transcriptomes of freshly isolated primary (1°) adult human AEC2s, their isogenic cultured progeny, and human iPSC-derived AEC2s (iAEC2s) cultured in identical conditions. We find each population occupies a distinct transcriptomic space with both types of cultured AEC2s (1° and iAEC2s) exhibiting similarities to and differences from freshly purified 1° cells. Across each cell type, we find an inverse relationship between proliferative states and AEC2 maturation states, with uncultured 1° AEC2s being most quiescent and mature, their cultured progeny being more proliferative/less mature, and cultured iAEC2s being most proliferative/least mature. iAEC2s also express significantly lower levels of major histocompatibility complex (MHC) genes compared to 1° cells, suggesting immunological immaturity. Cultures of either type of human AEC2 (1° or iAEC2) do not generate detectable type 1 alveolar cells in these defined conditions; however, iAEC2s after co-culture with fibroblasts can give rise to a subset of cells expressing “transitional cell markers” recently described in fibrotic lung tissue of patients with pulmonary fibrosis or in mouse models of pulmonary fibrosis. Hence, we provide direct comparisons of the transcriptomic programs of 1° and engineered AEC2s, two *in vitro* model systems that can be harnessed for studies of human lung health and disease.

## Introduction

Lung alveolar epithelial type 2 cells (AEC2s) fulfill specialized functions in their quiescent state and serve as facultative progenitors, able to re-enter cell cycle in order to maintain the alveolar epithelium after injury. Dysfunction of this key cell type has been implicated in the pathogenesis of a number of pulmonary diseases, including pulmonary fibrosis (1–4). Limited access to primary (1°) AEC2s from patients and difficulties with their phenotypic maintenance in long term *ex vivo* cultures has impeded development of tractable *in vitro* disease models. Prior investigations have focused on the development of methods for purification and *in vitro* propagation of rodent or human AEC2s, initially using 2D culture methods (5–7) and more recently using 3D culture conditions with (8–12) or without (13–16) supporting feeder cells. Early attempts to maintain AEC2s in 2D culture were limited by rapid loss of AEC2 specific gene expression and loss of proliferative capacity (8, 17), with several investigators noting the upregulation of gene markers interpreted by some (5, 17–19), but not all (20), as indicating differentiation of cuboidal AEC2s into squamous appearing “alveolar epithelial type 1 (AEC1)-like cells”.

With the advent of 3D culture models, we and others have developed methods for the *in vitro* culture of human AEC2-like cells derived from either 1° fetal (7), 1° adult (10, 14–16, 21, 22), or induced pluripotent stem cell (iPSC) sources (13, 22, 23). These methods have enabled the *in vitro* maintenance of functional human AEC2-like cells that share similar transcriptional and ultra-structural properties to their *in vivo* or freshly isolated 1° AEC2 counterparts, including the capacity to produce surfactant proteins and phospholipids (13–16, 23). Despite these advances which now enable more prolonged maintenance in cell culture of human AEC2-like cells, many controversies and questions remain to be addressed. For example, 1° or iPSC-derived AEC2-like cells captured in cell culture have been compared to *in vivo* 1° cells by bulk methods, such as RT-qPCR (23) or RNA sequencing (13, 23), but have yet to be directly compared head-to-head (without potential technical batch effects) at the single cell level to their uncultured 1° cell counterparts, raising uncertainty regarding how closely either 1° cell-derived or iPSC-derived AEC2-like cells resemble *in vivo* AEC2 controls, in terms of global gene expression profiles, heterogeneity, and proliferation states. In addition, the differentiation repertoire of human AEC2s either *in vivo* or *in vitro* remains uncertain. Few studies have provided comprehensive profiles, beyond just a few selected markers of unclear specificity, of human AEC1s arising from human AEC2s in culture (reviewed in (22)). Furthermore, some studies (10) found no evidence of AEC1s arising from 1° adult human AEC2s when cultured as alveolospheres. These findings for cultured human AEC2 cells are in contrast with observations made in mice, where lineage tracing studies as well as single cell RNA sequencing (scRNA-seq) profiles have established that adult rat or mouse AEC2s give rise to AEC1s both *in vitro* and *in vivo* (10, 24–27). In addition, a rapidly emerging literature has revealed a variety of transitional lung epithelial states appearing in cultured mouse AEC2 samples, in mouse lung injury models, or in the distal lung tissues of patients with fibrosing illnesses (24, 25, 27–30). These transitional cells have been identified by a diversity of markers not expressed in normal AEC2s, such as basal cell-like cytokeratins. Although neither the pathogenic/reparative potential of these transitional cells nor their cellular origins have been clearly determined, some have proposed AEC2s as their source (24, 25, 27, 31, 32); thus, whether analogous cells can be generated for study *in vitro* from human AEC2s is of increasing interest to those studying pulmonary fibrosis.

Here, we perform head-to-head comparisons between the single cell transcriptomes of 1° adult human AEC2s (prior to culturing), their isogenic cultured progeny, and human iPSC-derived AEC2s (iAEC2s) cultured in identical defined medium with and without feeders. We find that each population (1° preculture, 1° cultured, and iPSC-derived) expresses a distinct transcriptomic profile. Both cultured cell populations exhibit maintenance of the AEC2-like phenotype *in vitro*, with gene expression similarities with freshly captured adult 1° AEC2s. We provide quantitative correlation scores comparing each 1° and cultured AEC2 population and we observe a gradient of cell cycle states and maturation gene expression levels that distinguishes each cell preparation. Neither population of cultured AEC2s shows evidence of differentiation towards bona fide AEC1s when cultured in these conditions, but a subset of cells when cocultured with mesenchymal cells can be stimulated to upregulate markers of transitional cell phenotypes reminiscent of the recently described transitional cell states observed in mouse injury model systems (24, 25, 31) and in human lung diseases such as idiopathic pulmonary fibrosis (IPF) (25, 31).

## Results

### Establishment of synchronous primary (1°) AEC2 and iAEC2 cultures

To perform head-to-head comparisons between the global transcriptomes of 1° AEC2s, cultured 1° AEC2s, and cultured iAEC2s, we sought to establish synchronous cultures with similar conditions that would allow control of potential batch and media effects. Distal lung preparations from adult donor lung explants from two individuals (PL1, primary lung 1; PL2, primary lung 2) were cryopreserved using methods we recently described ((21) and Figure 1A). After thawing, AEC2s were purified using fluorescence activated cell sorting (FACS) to isolate cells co-expressing EPCAM and the AEC2 cell-selective surface marker HTII-280 (33). Once sorted, these cells were combined with MRC5 fibroblasts on cell culture inserts and *in vitro* colony forming efficiencies (CFE) were scored after outgrowth in three different media: CK+DCI, a defined serum-free medium that we have previously published for maintenance of human iAEC2s (13), “3D” medium previously published for the co-culture of mouse AEC2s with mesenchymal cells (11), or small airway epithelial cell growth medium (SAGM) (12) (Figure 1A, B). CK+DCI medium resulted in significantly higher CFE than the other 2 media and was thus chosen for further studies comparing cultured 1° AEC2s vs. iAEC2s in identical medium. Primary AEC2s cultured in the absence of supporting MRC5 fibroblasts did not yield any outgrowth colonies in these media (data not shown), consistent with our prior description of the need for supporting cells in order to culture 1° AEC2s (13). To test the expansion potential of 1° AEC2s co-cultured with MRC5 fibroblasts, we monitored growth kinetics and expression of the AEC2 program after serial passaging of each sample. After 1 passage (P1), 1° cultured AEC2s demonstrated significantly reduced CFE compared to the starting passage (P0) (Figure 1C) and displayed reduced expression of some AEC2-specific transcripts (e.g. *SFTPC*), but maintenance of others (e.g. *NKX2-1* and *SLC34A2*), when compared to pre-cultured and P0 1° AEC2s (Figure 1D). Primary AEC2s from neither of the tissue donors could be propagated beyond 2 serial passages (Figure 1C), whereas iAEC2s could be passaged indefinitely with no requirement for co-cultured fibroblasts, as previously published (34, 35) (Figure 1C). iAEC2s expressed *SFTPC* at similar levels, *NKX2-1* at higher levels, and *SLC34A2* at lower levels compared to 1° P1 AEC2s (Figure 1D).

**Figure 1.**
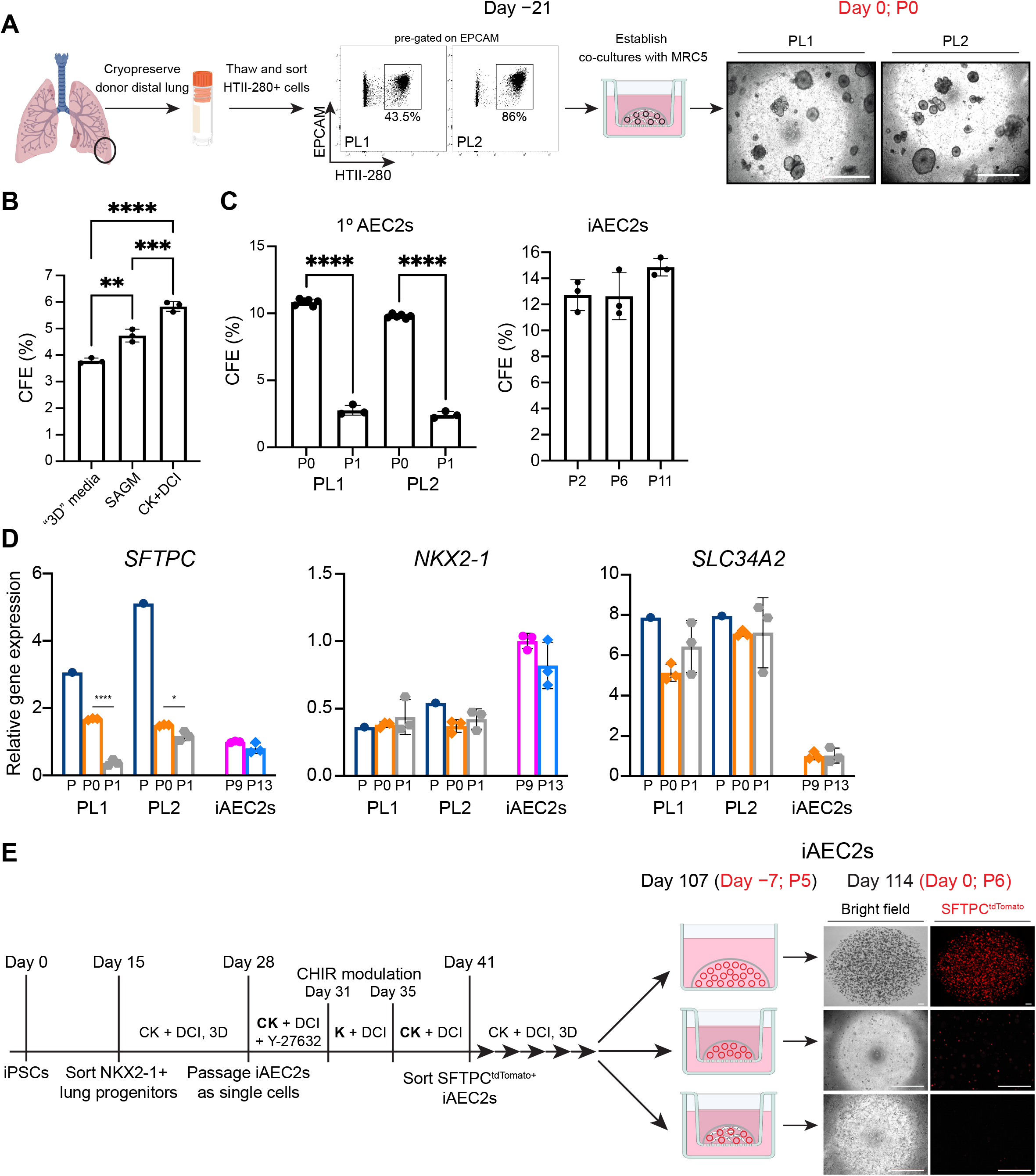
Establishment of synchronous primary (1o) AEC2 and iAEC2 cultures. (A) Schematic depicting the cryopreservation of distal lung preparations from adult donor lung explants (PL1, primary lung 1; PL2, primary lung 2), flow cytometry dot plots with gates used to isolate cells co-expressing EPCAM and the AEC2 cell-selective surface marker HTII-280 which were then combined with MRC5 fibroblasts on cell culture inserts on day −21. Representative live-cell imaging of the outgrowths on the day of encapsulation for scRNA-seq (day 0). Scale bars: 500 μm. (B) Bar graph shows the colony forming efficiencies (CFE) of 1o AEC2s in three different media. Bar graphs showing CFE after the first plating in culture prior to passaging (P0), reduced CFE of passaged (P1) 1o AEC2s, and stable CFE of iAEC2s across multiple passages. Notably, no colonies were formed from P2 1o AEC2s. (D) RT-qPCR showing fold change in gene expression compared to P9 iAEC2s in P13 iAEC2s and pre-culture (P) and cultured (P0 and P1) 1o AEC2s from two different donors (PL1 and PL2). (E) Schematic of directed differentiation protocol from iPSCs to day 107 (P7) monolayered epithelial iAEC2 spheres (alveolospheres). Seven days prior to encapsulation for scRNA-seq (day −7), 3D iAEC2s were dissociated and plated in 3 parallel conditions: 1) continued 3D feeder-free iAEC2 cultures, 2) feeder-free cultures on cell culture inserts, or 3) insert cultures with MRC5 fibroblasts identical to conditions for the 1o AEC2s. Representative live-cell imaging of the outgrowths on the day of encapsulation for scRNA-seq (day 0). Scale bars: 500 μm. (B-D) Mean ± SD is shown; n=3 experimental replicates). *p<0.05, **p<0.01, ***p<0.001, ****p<0.0001 by unpaired, two-tailed Student’s t-test for all panels.

To establish synchronous cultures of 1o AEC2s and iAEC2s for profiling of single cell transcriptomes, we used our previously published lung directed differentiation protocol (13, 36) to establish 3D cultures of indefinitely self-renewing pure iAEC2s from a human iPSC line carrying a tdTomato fluorescent reporter targeted to the SFTPC locus (SPC2 line; SPC2-ST-B2 clone) (34, 35) (Figure 1E). In these feeder-free 3D culture conditions, iAEC2s demonstrate maintenance of an AEC2 specific transcriptomic profile (Figure 1D; see also (35)). In parallel wells, we dissociated 3D iAEC2s and plated their progeny in CK+DCI medium in 3 parallel conditions: 1) continued 3D feeder-free iAEC2 cultures, 2) feeder-free cultures on cell culture inserts, or 3) insert cultures with MRC5 fibroblasts (Figure 1E) that were identical to conditions for the 1o AEC2s, cultured in parallel. In contrast to iAEC2s maintained in 3D feeder-free conditions, co-culture of iAEC2s with MRC5 fibroblasts resulted in loss of SFTPCtdTomato expression (Figure 1E) suggesting that these fibroblasts alter the molecular phenotype of iAEC2s.

### Single-cell transcriptomic profiling of freshly-isolated primary (1o) AEC2s, cultured 1o AEC2s, and iAEC2s

Next, we used scRNA-seq to compare 7 samples, all prepared in parallel and sequenced on the same day in order to avoid technical batch effects. We profiled each of the above 3 iAEC2 cell preparations along with 4 different 1o AEC2 samples consisting of: (a) “pre-culture 1o AEC2s”, defined as freshly isolated, cryopreserved HTII-280+ 1o AEC2s FACS-purified from the same two donor lungs used to establish the above 1o AEC2 cultures (Figure 1A), and (b) “cultured 1o AEC2s”, defined as the P0 1o AEC2 progeny of the “pre-culture 1o AEC2s” after 21 days of culturing in CK+DCI medium with MRC5 fibroblasts on cell culture inserts. The iAEC2 sample co-cultured with MRC5s on inserts as well as the two cultured 1o AEC2 samples were sorted for viable EPCAM+ cells prior to scRNA-seq. All other samples were sorted for live cells only. In order to harvest cells of similar confluence despite different outgrowth kinetics of 1o AEC2s vs iAEC2s, iAEC2s were harvested on day 7 after passage, whereas P0 1o AEC2s were harvested on day 21. We found that the transcriptomes of each AEC2 population (1o pre-culture, 1o cultured, and iPSC-derived) occupied distinct transcriptomic spaces when visualized by uniform manifold approximation and projection (UMAP; Figure 2A). Louvain clustering identified 16 clusters of cells (Figure 2B) driven primarily by sample type or donor identity, but with sub-clustering within cultured cell populations suggesting the presence of cell heterogeneity in vitro.

**Figure 2.**
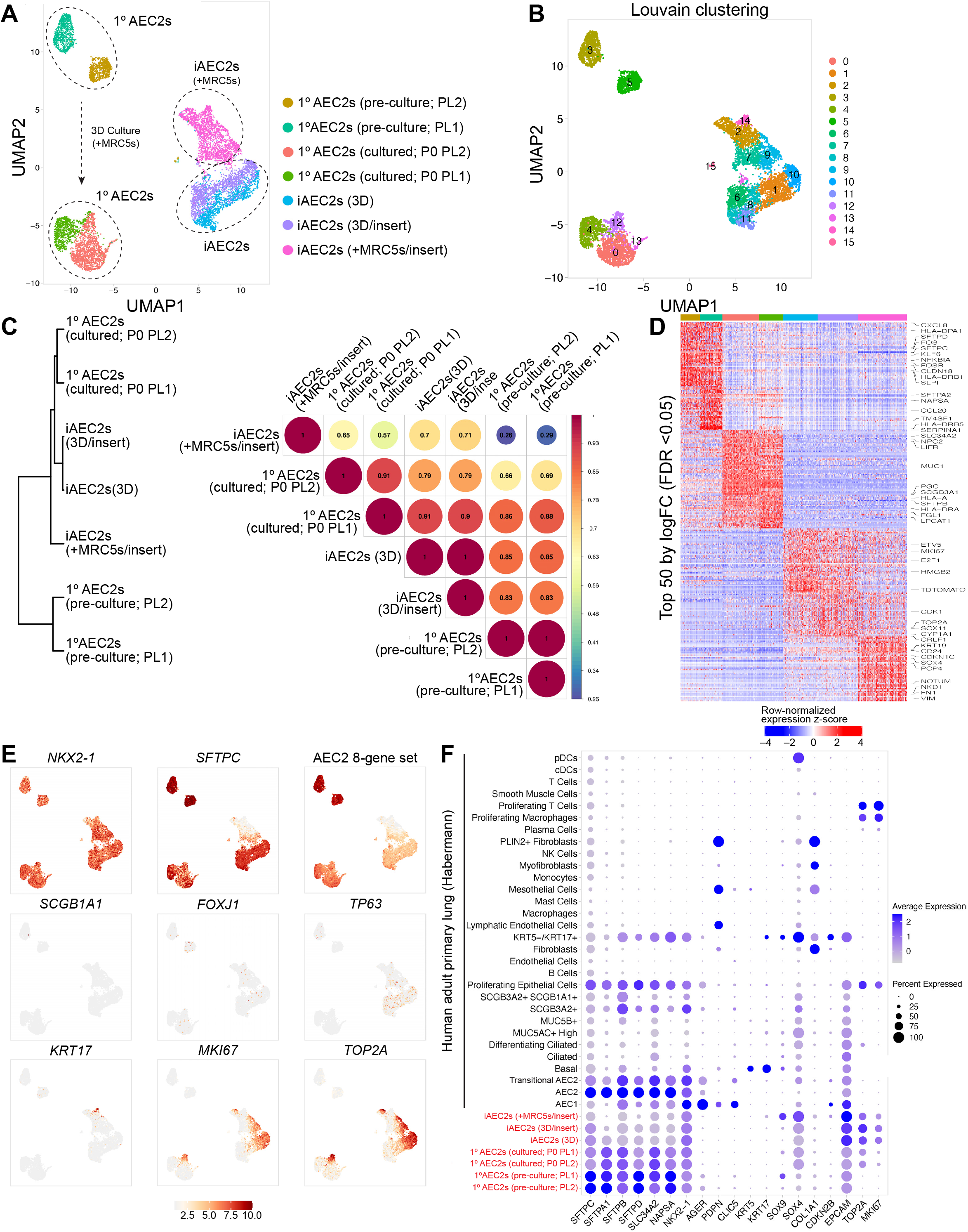
Single-cell transcriptomic profiling of 1o AEC2s and iAEC2s. (A) Visualization of pre-culture 1o AEC2s, cultured 1o AEC2s, and iAEC2s scRNA-seq transcriptomes using Uniform Manifold Approximation Projection (UMAP). (B) Louvain clustering of cell transcriptomes identifies 16 different clusters of cells driven primarily by sample type or donor identity, but with sub-clustering within cultured cell populations suggesting the presence of cell heterogeneity in vitro. (C) Dendrogram and heatmap of Pearson correlation coefficients between each sample based on normalized expression of the 3000 most variable genes across all cells. (D) Heatmap of top 50 differentially upregulated genes for each sample by scRNA-seq (ranked by average log fold-change, FDR <0.05; row-normalized expression z-scores). A subset of differentially expressed genes is highlighted with large font. (E) Normalized gene expression overlayed on UMAP plots for the indicated transcripts or gene sets. (F) Average expression levels and frequencies (purple dots) for select genes profiled by scRNA-seq in pre-culture 1o AEC2s, cultured 1o AEC2s, and iAEC2s. Comparison is made to a publicly available adult 1o distal lung dataset (30) and genes are selected to indicate AEC2, AEC1, airway, endothelial, epithelial, leukocyte, or proliferation programs.

To quantitatively score similarities in gene expression between each sample, we plotted the normalized expression of the 3000 most variable genes across all cells and calculated Pearson correlation coefficients for pairwise sample comparisons (Figure 2C). We observed high gene expression correlations between each sample with the exception of MRC5 co-cultured iAEC2s. Specifically, precultured vs. cultured 1o AEC2s displayed high correlations, as did feeder-free iAEC2s vs. pre-culture 1o AEC2s, and feeder-free iAEC2 vs. cultured 1o AEC2s. In contrast, iAEC2s after co-culturing with MRC5s displayed lower correlations with all other samples (Figure 2C). Together, these data indicate that cultured 1o AEC2s and cultured feeder-free iAEC2s exhibit transcriptomic similarities to as well as differences from pre-culture 1o AEC2s.

Focusing first on the canonical AEC2 marker, SFTPC, as anticipated by bulk RT-qPCR profiles (Figure 1D), we observed significantly higher expression of this transcript in pre-cultured cells by scRNA-seq, although expression was still easily detected in the vast majority of 1o AEC2s or iAEC2s after culturing (Figure 2D-F). In contrast, after co-culturing with MRC5 fibroblasts, iAEC2s expressed less SFTPC (Figure 2D, E), as predicted based on loss of SFTPCtdTomato reporter gene expression (Figure 1E) and also lower NKX2-1 (Figure 2D). This loss of SFTPC in co-cultured iAEC2s was consistent with their lower Pearson correlation scores compared to other cells, (Figure 2C). In addition to loss of SFTPC in this sample, a distinct cell cluster (cluster 14; Figure 2B) enriched in cytokeratins (KRT8, KRT17, KRT19) emerged (Figure 2D-F and further discussed below). Importantly, cultured 1o or iPSC-derived cells did not detectably assume alternative proximal or distal lung fates based on little to no expression of airway (SCGB1A1, FOXJ1, TP63, KRT5; Figure 2E, F) or AEC1 transcripts (Figure 2F).

Next, we assessed the most significant differences in gene expression between samples. Among the top 50 differentially upregulated genes in pre-cultured 1o AEC2s compared to all other cells (Figure 2D) were transcripts associated with AEC2 differentiation or maturation (34) (SFTPA2, SFTPC, SFTPD, NAPSA, SLPI) and immune related transcripts (CXCL1, CXCL2, CXCL3, CXCL8, CCL2, CCL20, HLA-DPA1, HLA-DRB1, HLA-DRB5, NFKBIA, NFKBIZ, TNFRSF12A, TNFAIP3, CD83). Cultured 1o AEC2s were also enriched in transcripts encoding AEC2-marker genes although their expression levels were lower than their pre-culture counterparts. iAEC2s were more proliferative than cultured 1o AEC2s based on significantly higher expression of MKI67, TOP2A, and transcripts associated with cytokinesis (Figure 2D-F). iAEC2s expressed the canonical AEC2 marker transcript SFTPC and the transcript for the SFTPCtdTomato reporter (Figure 2D-F). Compared to all other cells, iAEC2s cultured in feeder-free conditions expressed significantly higher levels of some distal alveolar epithelial marker transcripts (ETV5, CRLF1; Figure 2D) and lower levels of most mature AEC2 marker transcripts (SFTPA2, SFTPD, NAPSA, SLPI).

As prior studies have observed an inverse relationship between the proliferation and maturation programs in AEC2s (13, 37, 38), we next focused on comparing the proliferation and maturation transcriptomic programs across all 7 samples. We found a continuum of progressively more proliferative states (from fresh 1o to cultured 1o to cultured iAEC2s) that were inversely associated with gene signatures of AEC2 maturation (Figure 3A-C). Specifically, scRNA-seq data confirmed that cultured AEC2s (1o or iAEC2s) were more proliferative compared to preculture 1o AEC2s, as expected. While 66.5% of iAEC2s and 18% of cultured 1o AEC2s expressed transcripts associated with active cell cycling (S, G2, or M phase), only 0.1% of preculture 1o AEC2s expressed such transcripts (Figure 3D).

**Figure 3.**
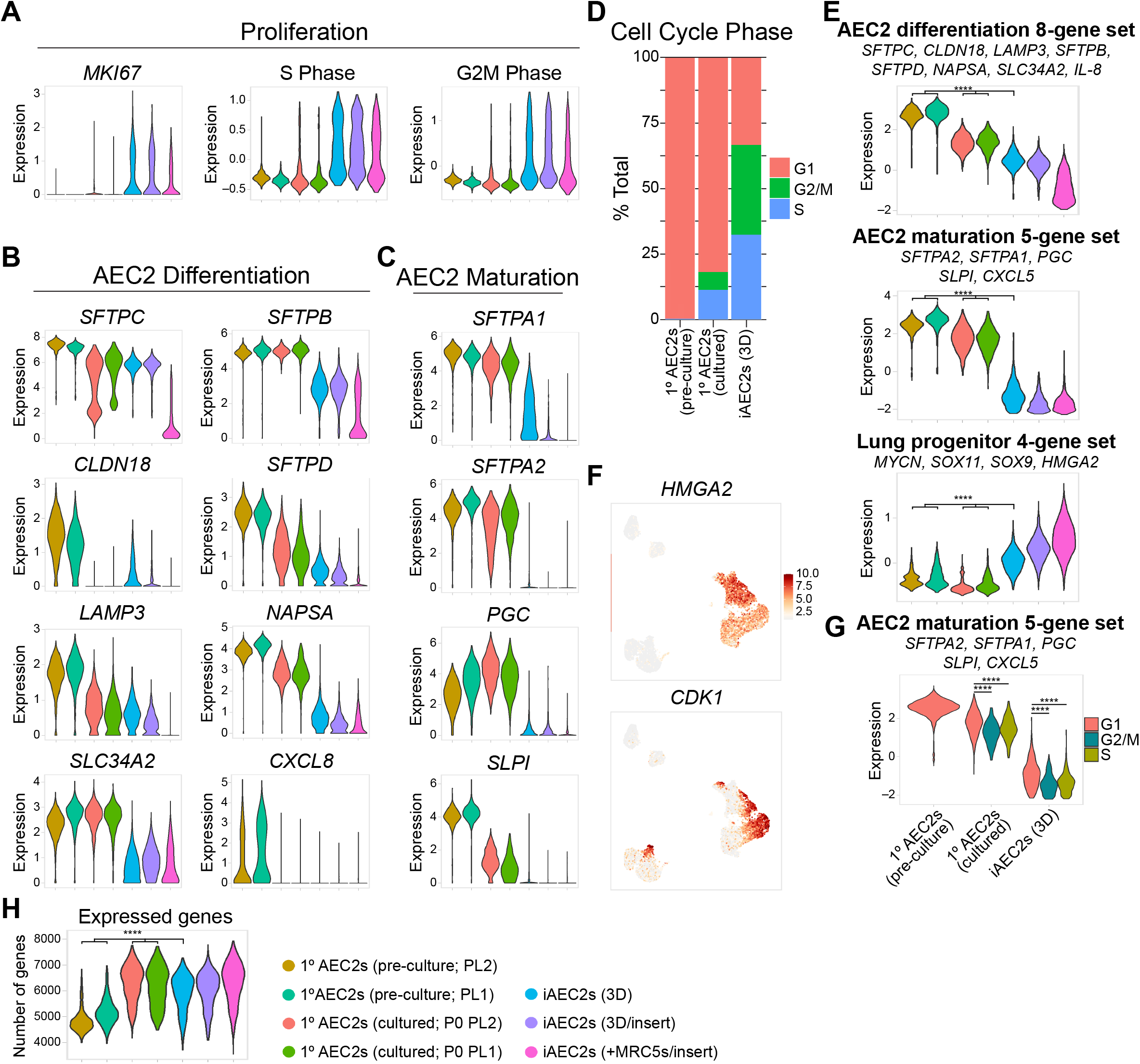
AEC2 maturation is inversely related to proliferation. (A) Violin plots showing normalized expression for MKI67 and cell cycle phase in pre-culture 1o AEC2s, cultured 1o AEC2s, and iAEC2s by scRNA-seq. (B) Violin plots showing normalized expression for individual genes from a published AEC2 differentiation gene set (34). (C) Violin plots showing normalized expression for individual genes from a published maturation gene set (34). (D) Barplot of cell cycle phase proportions by sample. (E) Violin plots showing normalized expression for indicated gene sets in pre-culture 1o AEC2s, cultured 1o AEC2s, and iAEC2s by scRNA-seq. (F) Normalized gene expression overlayed on UMAP plots for the indicated transcripts. (G) Violin plots of AEC2 maturation gene set by cell cycle phase in pre-culture 1o AEC2s, cultured 1o AEC2s, and iAEC2s. (H) Violin plots of expressed genes in pre-culture 1o AEC2s, cultured 1o AEC2s, and iAEC2s by scRNA-seq. ****p<0.0001 by unpaired, two-tailed Student’s t-test for all panels.

In contrast, top genes upregulated in pre-culture 1o AEC2s included transcripts encoding surfactants, lamellar body-related, and other AEC2-marker genes (Figure 2D-F). Similarly, pre-culture 1o AEC2s expressed higher levels of genes and gene sets (34) associated with AEC2 differentiation (Figure 3B, E) and maturation (Figure 3C, E) whereas cultured AEC2s expressed higher levels of genes (HMGA2, CDK1) and gene sets (MYCN, SOX11, SOX9, HMGA2; (34)) associated with progenitor cells (Figure 3E, F). Across the different samples, cells that were not in S/G2/M phases expressed significantly higher levels of the AEC2 maturation gene set, again suggesting an inverse relationship between AEC2 maturation and proliferation (Figure 3G). Lastly, cultured AEC2s expressed a higher number of genes when compared to preculture 1o AEC2 (Figure 3H).

Comparing feeder free iAEC2s to cultured 1o AEC2s, we found the distal lung progenitor marker, HMGA2, as well as Wnt target gene WIF1, to be in the top 25 transcripts differentially upregulated in iAEC2s (Figure 4A). In addition, consistent with their faster growth kinetics and greater passageability (Figure 1), iAEC2s compared to cultured 1o AEC2s expressed significantly more MKI67, TOP2A, and gene sets associated with active (S phase or G2M phase) cell cycle (Figure 4B). Consistent with this paradigm and the observation that 1o AEC2s are less proliferative than iAEC2s, we found markers of AEC2 differentiation and maturation (SFTPA2, NAPSA, SLC34A2, and ABCA3) to be in the top 25 most differentially upregulated transcripts in cultured 1o AEC2s compared to feeder-free iAEC2s. Gene set enrichment analysis (GSEA) indicated p53 signaling among the top upregulated pathways in cultured 1o AEC2s vs iAEC2s and cell cycle pathways (E2F, MYC, G2M checkpoint signaling) among the top downregulated (Figure 4C), consistent with their less active cell cycle and less proliferative state than iAEC2s.

**Figure 4.**
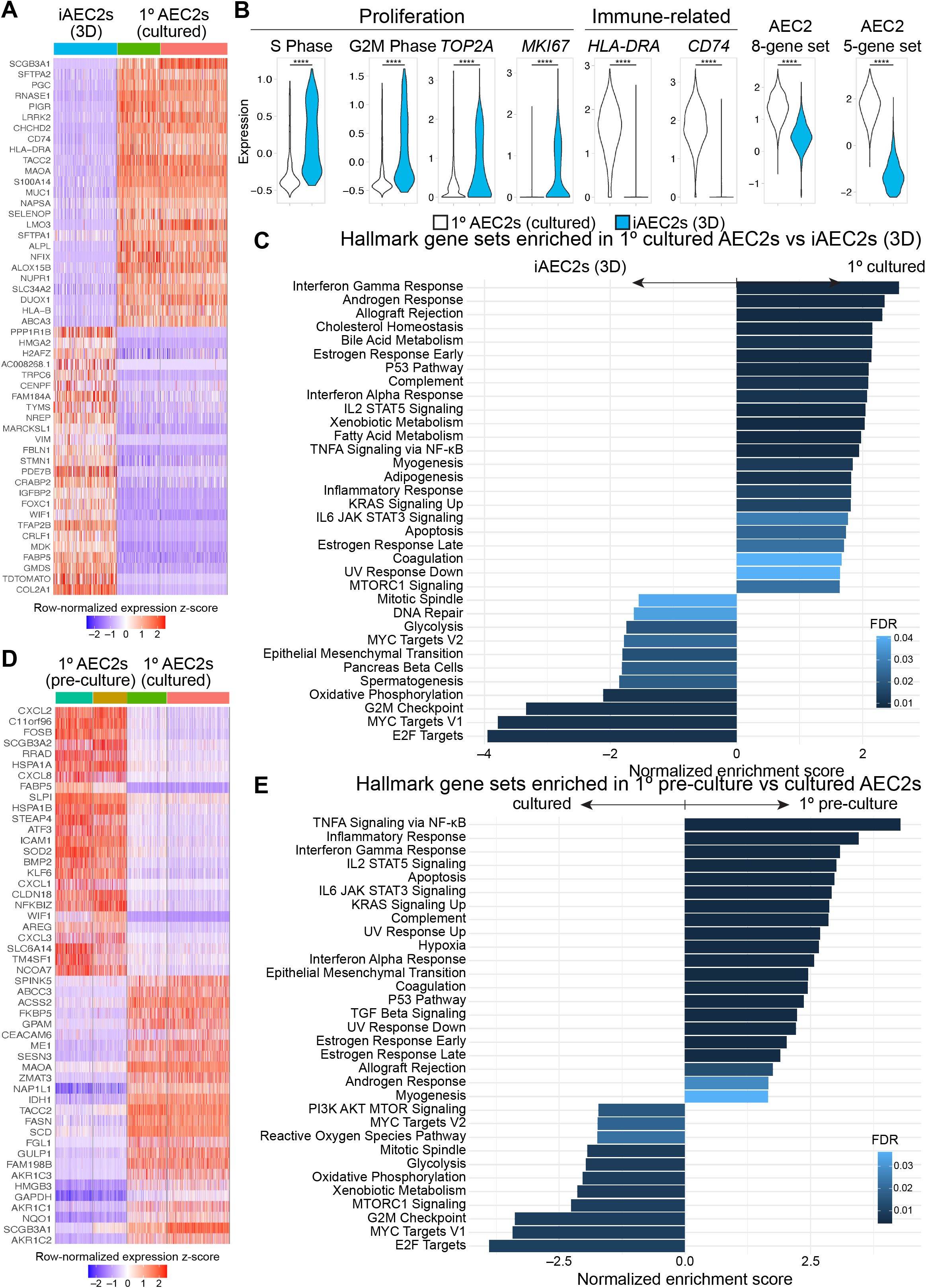
Pairwise single-cell transcriptomic comparisons of AEC2s. (A) Heatmap of top 25 downregulated and top 25 upregulated genes comparing feeder-free iAEC2s to cultured 1o AEC2s by scRNA-seq (ranked by average log fold-change, FDR <0.05; row-normalized expression z-scores). (B) Violin plots showing normalized expression for indicated genes, gene sets, or cell cycle phase in 1o cultured AEC2s vs iAEC2s by scRNA-seq. (C) Gene set enrichment analysis (GSEA, camera using Hallmark gene sets) of differentially regulated gene sets in cultured 1o AEC2s vs feeder-free (3D) iAEC2s (FDR <0.05). (D) Heatmap of top 25 downregulated and top 25 upregulated genes comparing cultured 1o AEC2s to pre-culture 1o AEC2s by scRNA-seq (ranked by average log fold-change, FDR <0.05; row-normalized expression z-scores). (E) Gene set enrichment analysis (GSEA, camera using Hallmark gene sets) of the top upregulated gene sets in cultured 1° AEC2s vs iAEC2s. (E) GSEA (camera using Hallmark gene sets) of differentially regulated gene sets in pre-cultured 1o AEC2s vs cultured AEC2s (cultured 1o AEC2s and feeder-free iAEC2s combined; FDR <0.05).

We performed further GSEA of cultured 1o AEC2s vs iAEC2s and found several additional pathways enriched in 1o cultured cells (Figure 4C). Multiple immune signaling pathways were more highly expressed in 1o cells, including NF-κB, interferon gamma, IL2/STAT5 signaling, and IL6/JAK/STAT3 signaling (Figure 4C). Most notably transcripts encoding MHC class I and II members (HLA-B, HLA-DRA, and CD74) were in the top 25 transcripts most downregulated in iAEC2s compared to cultured 1o AEC2s (Figure 4A, B), consistent with our prior published work (13) finding that iAEC2s express significantly less pathways and genes associated with immune maturation compared to 1o AEC2s.

Some loss of immune-related transcripts was a consistent feature across culture platforms, including in cultured 1o cells. For example, comparing pre-culture 1o AEC2s to their cultured (P0) progeny, we found a number of immune related transcripts (CXCL1, CXCL2, CXCL3, CXCL8, NFKBIZ) to be in the top 25 transcripts differentially downregulated with culture (Figure 4D). The Wnt target gene WIF1 as well as TM4SF1, a conserved cell surface marker of the previously described Wnt-responsive alveolar epithelial progenitors (12), were also among the top differentially downregulated transcripts after culturing 1o AEC2s (Figure 4D). Consistent with this, GSEA of pre-culture 1o vs all cultured AEC2s (1o AEC2s and iAEC2s combined) identified a number of immune related pathways as being significantly downregulated in cultured cells, whereas proliferation related pathways were enriched in cultured AEC2s (Figure 4E).

### Human AEC2s cultured in CK+DCI do not give rise to AEC1s

Next, we assessed whether there was emergence of AEC1s in our cultured AEC2s or iAEC2s. Notably there are few, if any, reports of entirely specific AEC1 marker genes whose expression in the human lung has been validated to unambiguously define the cell type. This contrasts with mouse lungs where a broad literature suggests several markers, including Hopx, that have been validated as being able to distinguish adult AEC1s from AEC2s (39–41). Thus, we selected multiple gene markers to screen for human AEC1s in our datasets. Using expression levels of a number of genes associated with AEC1s (PDPN, HOPX, AGER, AQP5, CAV1, CLIC5) (40, 42–46) (Figure 5A) or immunostaining for AEC1 markers such as RAGE (Figure 5B), we found no evidence that human 1o AEC2s or iAEC2s differentiate to yield bona fide AEC1s when cultured either alone or in the presence of stromal support cells. The presence of a few AEC1s (co-expressing all canonical AEC1 markers, including AGER, CLIC5, CAV1, EMP2, and PDPN) in the HTII-280 sorted pre-culture 1o AEC2 samples (Figure 5A) as well as RAGE positive AEC1s in immunostained control lung sections (Figure 5C) provided positive controls for expression levels of these markers and further supported our interpretation that there was little to no detectable expression of a comparable AEC1 program in any of our cultured 1o AEC2s or iAEC2s, which is in agreement with previous studies (10, 13). Taken together, these data suggest that none of the AEC2 culture models evaluated in this study yield cell types whose transcriptomes reflect mature human AEC1s.

**Figure 5.**
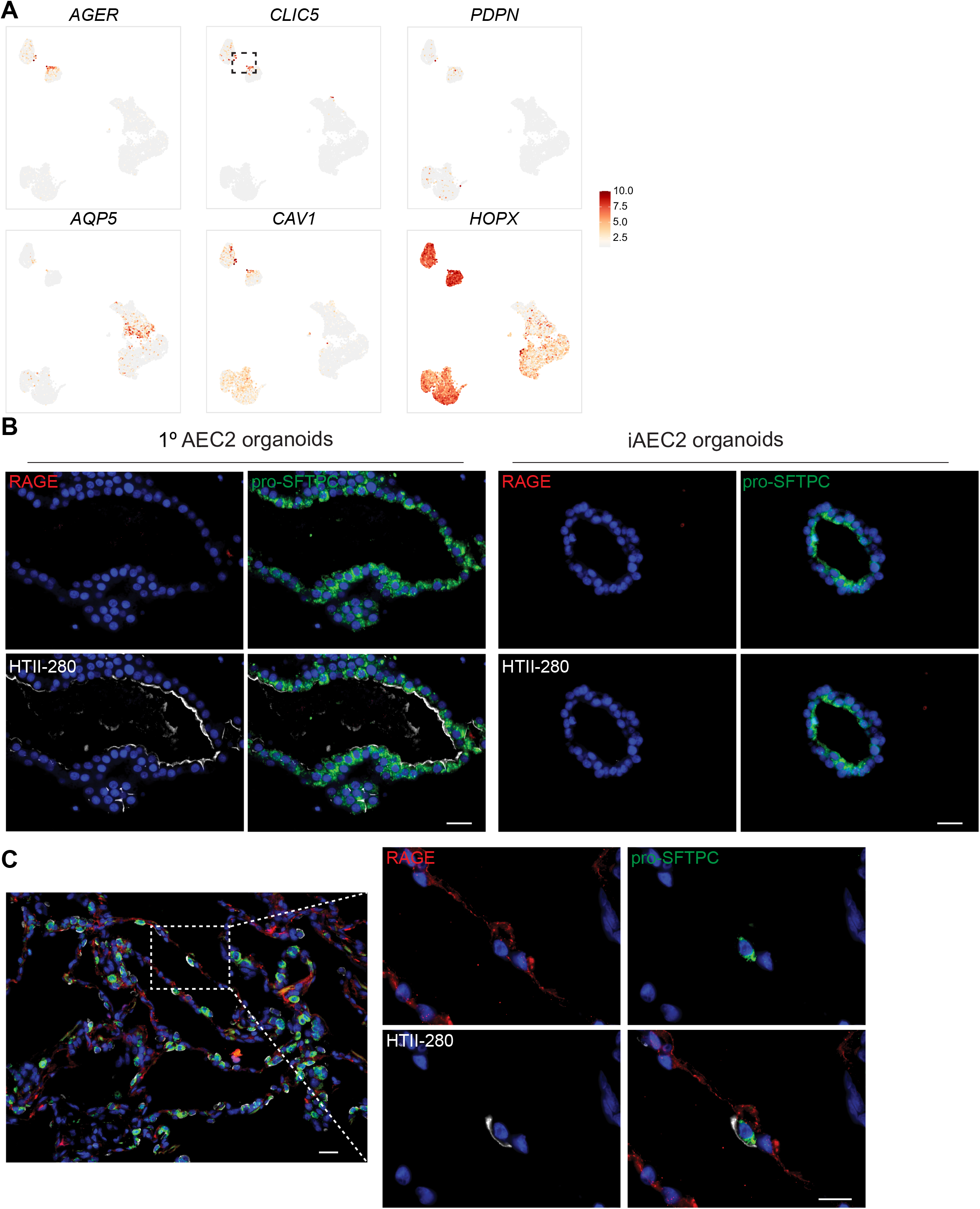
Absence of expression of the AEC1 molecular phenotype in cultured human AEC2s. (A) Normalized gene expression overlayed on UMAP plots for the indicated transcripts. Dashed gate shows the few 1o AEC1s. (B) Representative immunofluorescence microscopy of 1o AEC2 and iAEC2 organoids stained for RAGE (red), pro-SFTPC (green), HTII-280 (white), and DNA (Hoechst, blue). Scale bars: 25 μm. (C) Representative immunofluorescence microscopy of control adult human lung sections stained for the same markers shown in (B). Scale bars: 25 μm.

### Emergence of a transitional cell state

Recent studies have described a transitional cell epithelial state detected in injured distal lungs, characterized by expression of a variety of markers not normally expressed in AEC2s and implicated by some authors in the pathogenesis of IPF (24, 25, 31). We thus screened our various samples for the presence of any cells expressing these newly described “transitional markers”. We found a subset of iAEC2s after co-culturing with MRC5 fibroblasts (Figure 2B; cluster 14) was highly enriched for KRT17 (Figure 2E) and other markers (KRT8, KRT19, SOX4, CLDN4) described in this transitional cell state (Figure 6A, B). Nine of the top 50 differentially upregulated genes in this cluster have previously been associated with a human KRT5-/KRT17+ transitional epithelial cell cluster identified in lungs of patients with end-stage IPF (30) (Figure 6B). iAEC2-derived cells expressing the KRT5-/KRT17+ gene set demonstrated lower expression of both SFTPC and members of an 8 gene signature that defines AEC2s (34). These cells did not express MKI67, suggesting they were quiescent (Figure 6C). To better map the origin of KRT5-/KRT17+ epithelial cells and since these cells presumably emerged from the starting iAEC2 population at some point over 7 days of co-culture with MRC5s, we performed RNA velocity analysis which was consistent with the interpretation that iAEC2s give rise to the KRT5-/KRT17+ cluster of cells over time (Figure 6D). To assess whether these cells might fully transition to AEC1s at a later time point, we repeated cultures of iAEC2s alone vs iAEC2s cultured with supporting cells and evaluated their gene expression weekly for 2 weeks, compared to cultured 1o AEC2 controls. We found that expression of both KRT17 and KRT8 increased over time in cultures of iAEC2s, particularly those cultured with MRC5 cells (Figure 6E). Cultured iAEC2s in these conditions again did not exhibit evidence of AEC1 differentiation based on low or no expression of AGER, PDPN, or CAV1 over the 2-week culture period (Figure 6E). Immunofluorescence staining confirmed the presence of KRT17 expression at the protein level in a subset of epithelial cells after co-culturing of iAEC2s with MRC5 fibroblasts (Figure 6F). These data support the emergence of a sub-population of transitional/intermediate epithelial cell state in iAEC2/fibroblast co-cultures that resembles transitional KRT5-/KRT17+ cells described in IPF lung tissue by Habermann et al. (30).

**Figure 6.**
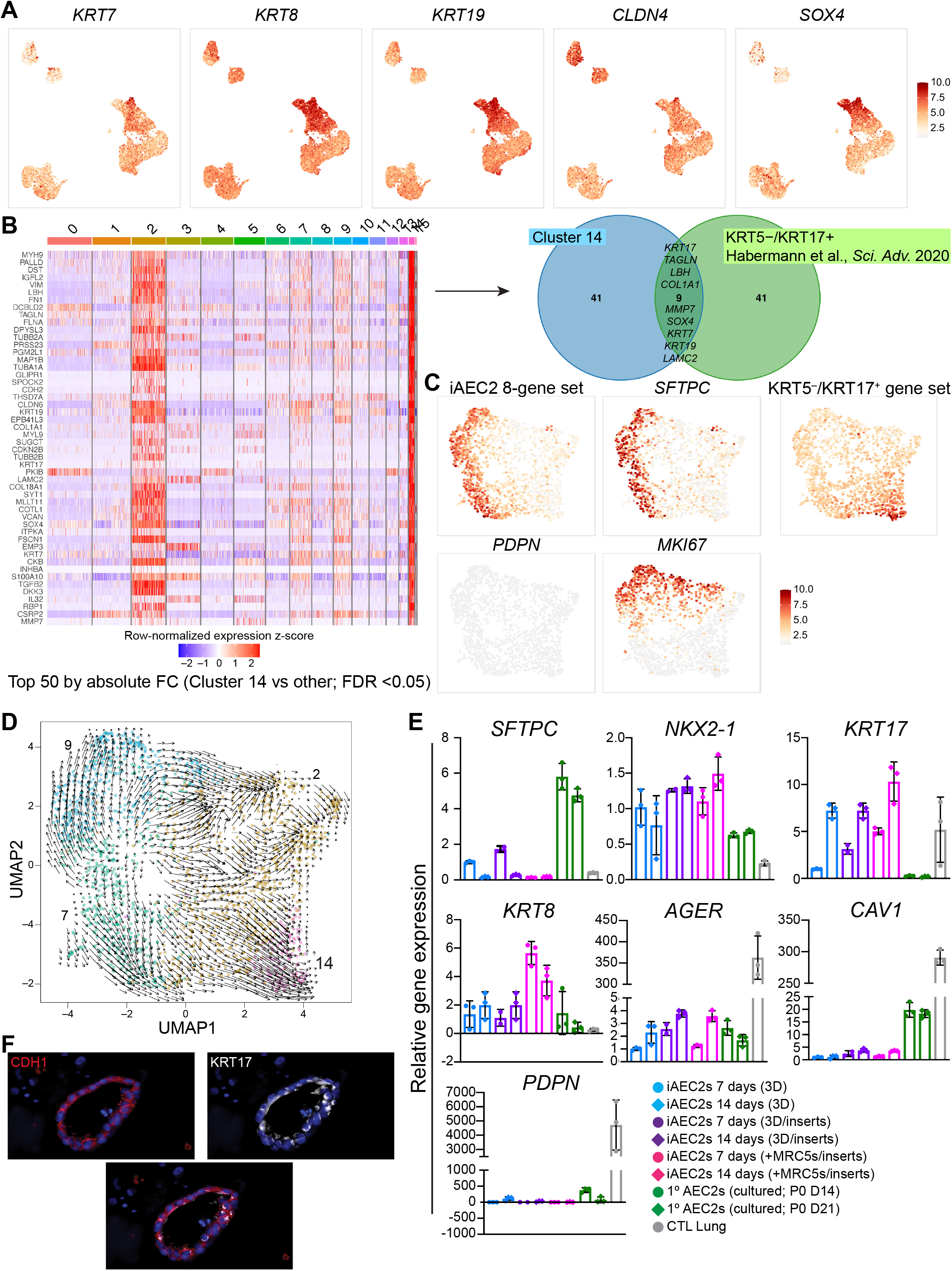
Emergence of a transitional cell state from iAEC2s co-cultured with MRC5 fibroblasts. (A) Normalized gene expression overlayed on UMAP plots for the indicated transcripts. (B) Heatmap of top 50 differentially upregulated genes comparing cluster 14 vs other clusters by scRNA-seq (ranked by average log fold-change, FDR <0.05; row-normalized expression z-scores). Venn diagram shows that 9 of the top 50 differentially upregulated genes in this cluster have previously been associated with a human KRT5-/KRT17+ transitional epithelial cell cluster (30). (C) Normalized gene expression overlayed on UMAP plots of the sample of iAEC2s cultured on inserts together with MRC5s. Indicated transcripts or gene sets are overlayed. (D) RNA velocity analysis indicates that cluster 14 cells (enriched in the KRT5-/KRT17+ gene set from (C)) arise from iAEC2s over time. (E) RT-qPCR showing fold change in gene expression compared to iAEC2s cultured in feeder-free conditions for 7 days in the indicated samples. (F) Representative immunofluorescence microscopy of iAEC2s co-cultured with MRC5 fibroblasts and stained for E-cadherin/CDH1 (red), KRT17 (white), and DNA (Hoechst, blue). Scale bars: 25 μm.

## Discussion

Our results herein suggest that cultured 1o AEC2s and iAEC2s maintain an AEC2-like phenotype in vitro and their transcriptomes at single cell resolution exhibit similarities to and important differences from 1o pre-culture AEC2s. Most notably, we find a continuum of progressively more proliferative states (from pre-culture 1o to cultured 1o to iAEC2s) correlating with greater passageability and inversely associated with AEC2 maturation state (Figure 4A-C). Furthermore, our results reveal that immune related programs such as those involving MHC genes, are expressed at progressively lower levels in cultured AEC2s (decreasing from preculture 1o AEC2s to cultured 1o AEC2s to iAEC2s) suggesting that in vivo AEC2s are immunologically more mature than cultured cells.

Importantly our results suggest that despite their limitations cultured 1o AEC2s and iAEC2s maintain key components of the human AEC2 program, and thus it is likely that both cell populations can serve as in vitro models of the human lung epithelium for disease modeling with relevant benefits or limitations that should be taken into account when selecting each type of model system. The different characteristics of 1o AEC2s (higher expression of immune-related transcripts) and iAEC2s (patient specificity, greater proliferation and passageability) indicate that each model can be applied in different settings. For example, cultured 1o AEC2s with their higher expression of immune related genes and pathways may serve as preclinical disease models to study the responses of the alveolar epithelium to infectious agents, such as severe acute respiratory syndrome coronavirus 2 (SARS-CoV-2) in situations where immune maturation may be particularly relevant (47). In cases where high levels of expression of MHC genes is not required, other human AEC2-specific responses to infectious agents, including SARS-CoV-2, may be studied in either cultured cell type since both 1o cultured AEC2s and iAEC2s express a wide variety of receptors required for viral entry, such as ACE2 and TMPRSS2 (47, 48). On the other hand, patient-specific iAEC2s can be used as a human preclinical disease model when unable to access 1o AEC2s at early disease stages or when the application of invasive procedures required to isolate adequate numbers of 1o AEC2s is prohibitive, such as in the case of severe interstitial lung disease (35). Patients with advanced lung disease are likely to have AEC2s with transcriptomic and epigenetic programs that are secondarily perturbed by drugs, infections, ventilators, or other tertiary insults that may be difficult to distinguish from earlier disease-initiating mechanisms. Since iAEC2s are derived from iPSCs generated by reprogramming, which erases the starting epigenome or disease state, patient-specific iAEC2s in most cases will have been “reset” and thus in theory the emergence of disease, potentially including relevant genetic and epigenetic AEC2 programs, may be replayed repeatedly, potentially from disease inception through late stages and drug responses (35). This sequence may be harder to study in cultured 1o AEC2s procured from some patients with advanced disease, particularly if the disease results in a paucity of AEC2s in vivo, or if residual epigenetic perturbations or lasting secondary effects are carried through in any cultured 1o AEC2s from those patients.

In terms of disease-relevant cells that may arise from parental AEC2s, the emergence of so-called transitional cells from the co-culture of iAEC2s with mesenchymal cells was an unexpected result with clinically-relevant implications. These iAEC2-derived transitional cells appear to express a transcriptomic phenotype overlapping with that described in the distal lung tissues of patients with fibrosing illnesses (24, 25, 28–31), suggesting that these culture conditions could be used to better understand the cellular origin of these cells and their potential pathogenic or reparative roles.

We found that cultured 1o AEC2s, compared to iAEC2s, express significantly higher levels of transcripts known to be expressed late or postnatally in AEC2 development, also known as AEC2 “maturation markers”. These markers include surfactant-encoding transcripts known to be expressed late in AEC2 development, such SFTPA1 and SFTPA2 (34). In contrast, early surfactants, SFTPC and SFTPB, are expressed at high levels in both cultured 1° AEC2s and iAEC2s. iAEC2s have already been successfully employed to study the processing of normal and mutant SFTPB and SFTPC proteins, including disease modeling using patient-specific mutant iAEC2s and their gene-edited progeny (13, 35); however, for studies focused on understanding the biology of late surfactants, such as SFTPA1 and SFTPA2, investigators may choose 1o AEC2s rather than iAEC2s, at least when using the culture conditions employed in this report.

Whether the expression of maturation markers can be further augmented in iAEC2s, to the levels we found expressed in 1o cells, is an area of ongoing research. For example, we recently reported that transitioning iAEC2s to 2D air-liquid interface culture conditions results in decreased proliferation and increased maturation, exemplified by augmented expression of both early and late surfactants (SFTPC, SFTPA1, SFTPA2 and secretion of tubular myelin) (48, 49). It remains to be seen, however, whether this level of augmented maturation is able to reach the high levels of expression exhibited by 1o AEC2s in the scRNA-seq profiles we present here.

A particular hurdle that has limited research using human AEC2s in the past has been the inability to stably expand 1o AEC2s in culture, which would be required for gene editing studies, high throughput drug screens, or future cell-based therapies. Our results indicate that iAEC2s provide a potential solution since they can be indefinitely expanded in 3D cell culture and exhibit a more proliferative state than 1o AEC2s either before or after culture. Indeed, we have previously shown that >1023 iAEC2s can be derived per input patient-specific iAEC2 via serial passaging over a 300-day culture period (35). Although 1o AEC2s could not be serially passaged extensively in our current culture conditions, since the preparation of this manuscript three reports have published methods for maintaining 1o human AEC2s in extensively self-renewing, feeder-free cultures (14–16). Future work will be required to compare fresh/uncultured 1° AEC2s to their progeny expanded in these new conditions as well as to iAEC2s to understand whether the transcriptomic programs and functional repertoire of AEC2s differs from the cells we have profiled in our conditions.

Many human iPSC-based model systems, such as those designed to derive cardiomyocytes or hepatocytes, are characterized by the presence of embryologically immature cells relative to their 1o control cell counterparts (50, 51). It remains unclear, however, whether the lower levels of maturation markers expressed in iAEC2s compared to 1o cells in our studies should be interpreted as a lack of developmental maturation vs the acquisition in culture of a highly proliferative state such as one that might occur in post-natal distal lung tissue in vivo in response to injury. Indeed, the use of enzymes to digest tissues or epithelial spheres into single cell suspensions for culture likely represents a significant injury to which AEC2s must respond if they are to survive and proliferate in our culture conditions. As in prior reports (37, 38), our profiles indicate an inverse relationship between AEC2 proliferation and maturation states. However, the lack of current in vivo benchmarking scRNA-seq datasets from human fetal developing and post-injury adult lungs limits a definitive determination of which in vivo stages or states (fetal/developmental or adult post-injury, for example) are most similar to either iAEC2s or 1o cultured AEC2s. Intense research efforts, such as LungMAP, that are designed to prepare these types of in vivo benchmarks should help to address this ongoing question.

Finally, a controversy raised by our findings is whether human AEC2s generate bona fide AEC1s in cell culture as has been extensively demonstrated with mouse AEC2s (10, 22, 24, 25). Despite recent studies (12, 14, 15, 47) suggesting human AEC2-to-AEC1 differentiation in vitro based on expression of a few selected markers, such as HTI-56 and additional bulk transcriptomic profiles of in vitro human alveolar cell derivatives (42, 52, 53), it remains uncertain how closely these cultured AEC1-like cells resemble their in vivo counterparts, if compared head-to-head. Gotoh and colleagues recently showed generation of AEC1-like cells from iAEC2s in feeder-dependent cultures (54). However, comprehensive profiling of our cultures at single cell resolution based on expression of a set of markers and confirmed by immunostaining, reveals little evidence of cultured adult human AEC2s giving rise to bona fide AEC1s, at least under our culture conditions, in agreement with some prior reports studying 1o or iPSC-derived human cells (10, 13).

In summary, our results suggest that culturing human AEC2s has defined and quantifiable impacts on their transcriptomic programs and proliferative states. By profiling these cells head-to-head and at single cell resolution we provide comparisons of fresh 1o AEC2s, cultured 1o AEC2s, and iAEC2s, revealing similarities and differences and suggesting that both cultured 1o AEC2s and iAEC2s can serve as in vitro models to study the role of human AEC2s in the pathobiology of the distal lung.

## Acknowledgements

The authors wish to thank all members of the Kotton, Stripp, and Kim Labs for insightful discussions. We thank Brian R. Tilton of the BUSM Flow Cytometry Core. We are grateful to Greg Miller and Marianne James of the Boston University Center for Regenerative Medicine (CReM) for maintenance and characterization of patient-specific iPSCs, supported by NIH grants NO1 75N92020C00005 and U01TR001810. The schematics in Figure 1 were created with BioRender.com. This work was supported by an I.M. Rosenzweig Junior Investigator Award from The Pulmonary Fibrosis Foundation and an Integrated Pilot Grant Award through Boston University Clinical & Translational Science Institute (1UL1TR001430) to K.D.A.; a Mexico in Harvard Foundation-Mexican Council of Science and Technology Fellowship to C.G.A.R.; NIH grant P01 HL108793 to B.R.S.; NIH grants R01 HL132266, R01 HL125821, R35HL150876, a Cystic Fibrosis Foundation Award KIM19P0, LONGFONDS | Accelerate, project BREATH, Gilda and Alfred Slifka, Gail and Adam Slifka, the Cystic Fibrosis/Multiple Sclerosis Fund Foundation Inc. and the Harvard Stem Cell Institute to C.F.K.; NIH grants U01HL148692, U01HL134745, U01HL134766 and R01HL095993 to D.N.K.; and an IDEAL Consortium Grant from Celgene/Bristol Myers Squibb to B.R.S., C.F.K., and D.N.K.

## Author Contributions

KDA, CGAR, PP, BRS, CFK, and DNK conceived the work. KDA, CGAR, PP, BRS, CFK, and DNK designed experiments. KDA, CGAR, CY, PP, JH, OTH, KM, BK conducted experiments and analyzed data. CVM and CY performed bioinformatics analysis. KDA, CGAR, PP, BRS, CFK and DNK prepared and edited the manuscript. All authors reviewed and approved the final version prior to submission.

